# A dormant sub-population expressing interleukin-1 receptor characterises anti-estrogen resistant ALDH+ breast cancer stem cells

**DOI:** 10.1101/821876

**Authors:** Aida Sarmiento-Castro, Eva Caamaño-Gutiérrez, Andrew H. Sims, Mark I. James, Angélica Santiago-Gómez, Rachel Eyre, Christopher Clark, Martha E. Brown, Michael D. Brooks, Daniel F. Hayes, Max S. Wicha, Sacha J. Howell, Robert B. Clarke, Bruno M. Simões

## Abstract

Estrogen receptor-positive (ER+) breast tumours are often treated with anti-estrogen (AE) therapies but frequently develop resistance. Cancer Stem Cells (CSCs) with high aldehyde dehydrogenase (ALDH) activity (ALDH+ cells) are reported to be enriched following AE treatment. Here we perform *in vitro* and *in vivo* functional CSC assays and gene expression analysis to characterise the ALDH+ population in AE resistant metastatic patient samples and an ER+ cell line. We show that the IL1β signalling pathway is activated in ALDH+ cells and data from single cells reveals that AE treatment selects for IL1R1-expressing ALDH+ cells. Importantly, we demonstrate that increased expression of *IL1R1* is observed in the tumours of patients treated with AE therapy and predicts for treatment failure. Single-cell gene expression analysis revealed that at least 2 sub-populations exist within the ALDH+ population, one proliferative and one quiescent. Following AE therapy, the quiescent ALDH+IL1R1+ population is expanded, which suggests CSC dormancy as an adaptive strategy that facilitates treatment resistance. Supporting this, analysis of AE resistant dormant tumours reveals significantly increased expression of *ALDH1A1*, *ALDH1A3* and *IL1R1* genes. Thus, we propose that targeting of ALDH+IL1R1+ cells will reverse AE resistance, including in patients with minimal residual disease.

## INTRODUCTION

Breast cancer (BC) represents 25% of all cancer diagnoses and is the fifth most common cause of death in women worldwide. Approximately 80% of BCs are positive for estrogen receptor expression (ER+ tumours) and are treated with anti-estrogen (AE) adjuvant therapies such as tamoxifen or fulvestrant. Despite the clear benefit of these drugs at reducing tumour recurrence, *de novo* or acquired resistance often occurs (Pan et al., 2017).

Cancer Stem Cells (CSCs) are a cellular population endowed with self-renewal properties, which are responsible for tumour progression and metastasis (Reya et al., 2001). Aldehyde Dehydrogenase (ALDH) activity is reported to be a CSC marker in human BC cells (Ginestier et al., 2007). ALDH+ cells are ER-negative, and likely to be resistant to the direct effects of AE therapy (Honeth et al., 2014). We have previously established that ALDH+ cells drive therapeutic resistance in ER+ BC tumours (Simoes et al., 2015).

Intra-tumour heterogeneity within BCs hinders accurate diagnosis and effective treatment. Understanding of the cellular diversity within the CSC population, especially at the single cell level, is limited. Given the importance of ALDH+ cells in promoting AE resistance, we investigated the gene expression pattern of this cellular population at the single cell level. This study reveals a previously uncharacterised level of heterogeneity within AE resistant CSCs, and identifies IL1R1 as a potential target in refractory and dormant BCs.

## RESULTS

### ALDH+ cells from AE-treated ER+ BCs have greater breast CSC activity than ALDH− cells

Previous research reported by our group (Simoes et al., 2015) established that AE treatment of BC Patient-Derived Xenograft (PDX) tumours in mice enriches for breast CSCs (BCSCs) with high ALDH enzymatic activity. To further investigate this AE resistant population, we isolated ALDH+ and ALDH− cells from 8 metastatic ER+ BCs undergoing AE therapies. There was significant inter-individual variation in the percentage of ALDH+ cells (range 0.32%-27.3%) (**Figures 1A and S1A**). Importantly, ALDH+ cells exhibited significantly greater BCSC activity as assessed by mammosphere formation than ALDH− cells in 7 out of 8 patient samples, and in 4 of these samples the mammosphere forming efficiency (MFE) was increased by more than 3-fold (**Figure 1B**). On average, ALDH+ cells from the 8 metastatic BC samples showed 3.8-fold greater MFE than ALDH− cells (p=0.001) (**Figure 1C**). Next, we investigated the *in vivo* tumour-initiating capabilities of ALDH+ cells isolated from the ER+ cell line MCF-7, following 6-day *in vitro* treatment with the AEs tamoxifen or fulvestrant (**Figure 1D**). Injection of 1,000 ALDH+ cells consistently gave rise to bigger tumours compared to the same number of ALDH− cells significantly so in tamoxifen and fulvestrant-treated cells (**Figure 1E**). This suggests a cytostatic effect of these drugs which acts specifically on the ALDH− cell population. Extreme Limiting Dilution Analysis (ELDA) revealed that on average the number of tumour-initiating cells was 4.2-fold higher in ALDH+ compared to the non-BCSC ALDH− cells in all three conditions tested (**Figure 1F**). As few as 100 ALDH+ cells gave rise to tumours in mice whereas 100 ALDH− cells failed to do so. These results highlight the increased tumour-initiating capabilities of the ALDH+ population in comparison to ALDH− cells, implying the need to characterize this population of CSCs that survive AE therapies.

**Figure 1.**
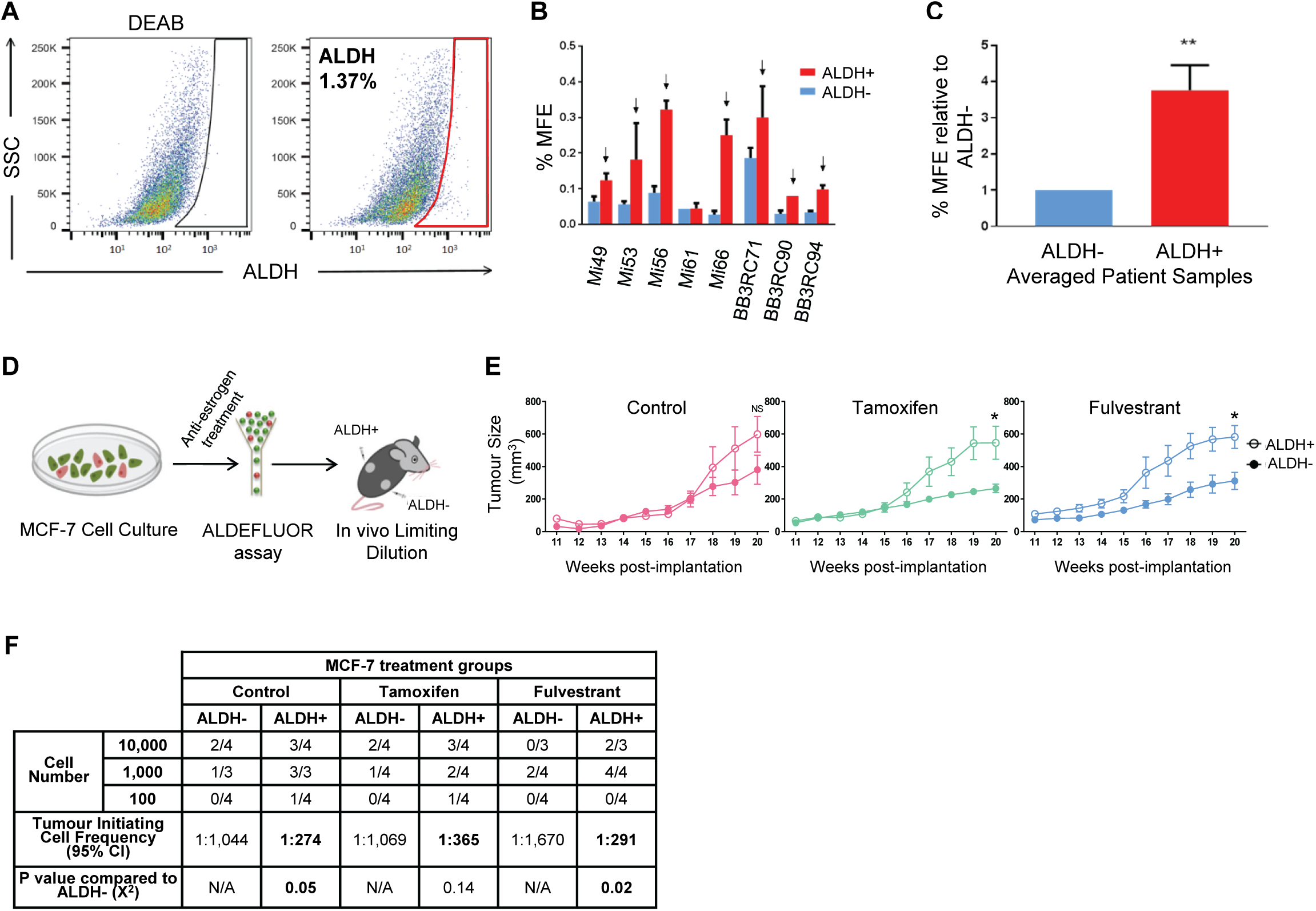
AE-treated ALDH+ cells from ER+ BC cells have greater BCSC activity than ALDH− cells *in vitro* and *in vivo*. (A) Representative FACS plot showing the ALDH+ population identified through the Aldefluor assay for an individual patient sample. ALDH+ cells (red gate) were discriminated from ALDH− cells using the DEAB control. (B) Bar chart shows mammosphere forming efficiency (MFE) percentage of ALDH+ cells (red) and ALDH− cells (blue) from ER+ metastatic BCs undergoing AE therapies. (C) Bar chart illustrates fold change in MFE percentage between ALDH+ and ALDH− cells across 8 different patient samples. (D) Schematic overview of the *in vivo* transplantation assay to test tumour formation capacity between ALDH+ and ALDH−MCF-7 cells. MCF-7 cells were pre-treated *in vitro* for 6 days with control (ethanol), tamoxifen (1 μM) or fulvestrant (0.1 μM) followed by the Aldefluor assay. ALDH+ and ALDH− cells were FACS sorted, counted using Trypan blue and then engrafted into the left and right flank, respectively, of the same NSG mice. (E) Averaged tumour growth from control (pink; left panel), tamoxifen (green; middle panel) or fulvestrant-treated (blue; right panel) cells. 1,000 ALDH+ (hollow circle) and 1,000 ALDH− (filled circle) cells are represented. *p≤0.05 (two tail, two sample equal variance t-test). N° of mice per condition =4 (vehicle-treated mice n=3). Data shows mean +/− SEM. (F) Table shows Extreme Limiting Dilution Analysis from *in vivo* injections of ALDH+ and ALDH− cells (10,000; 1,000; 100 cells) to assess tumour-initiating cell frequency. Tumor growth was assessed at week 20 and is represented as mice positive for growth/mice tested, for each cell number.

### Transcriptomic characterisation of ALDH+ cells in therapy resistant patient samples

To better understand the development of resistance to AE therapies in ER+ BC patients, we interrogated and compared the gene expression pattern between ALDH+ and ALDH− cells in 9 metastatic ER+ patient samples (**Figure 2A**). Overall, 599 genes were found to be differentially expressed (p≤0.05) between the two cell populations amongst the 18,752 genes with measured expression (**Table S1**).

**Figure 2.**
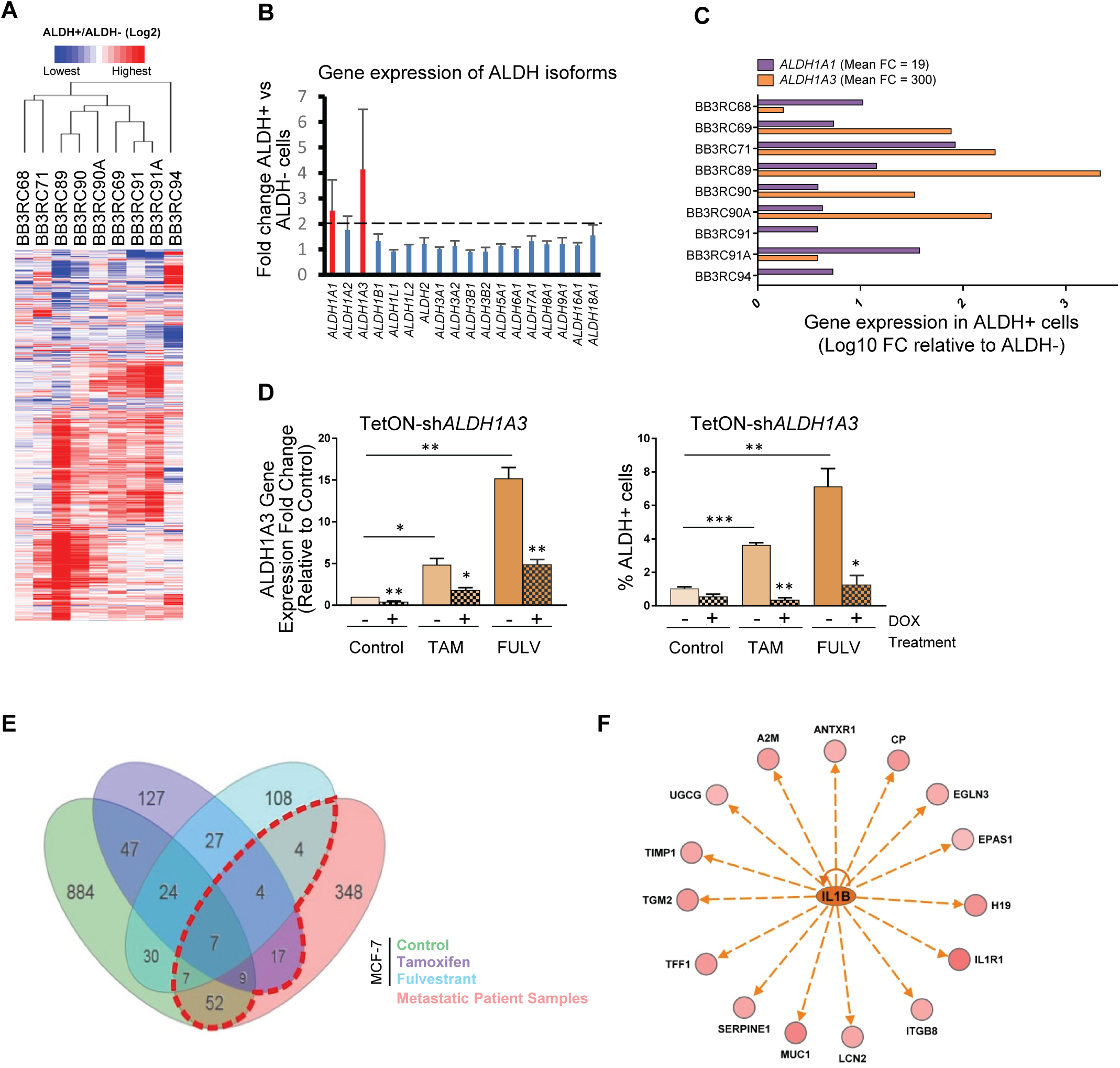
ALDH+ cells from AE resistant patient samples show a distinct gene expression pattern compared to ALDH− cells. (A) Heatmap illustrating the 599 differentially expressed genes (447 up, 152 down) between ALDH+ and ALDH− cells (red colour shows gene up-regulation, green shows down-regulation in ALDH+ relative to ALDH− cells identified by pairwise Rank Products with a threshold probability of false positives <0.05) from metastatic ER+ patient BCs undergoing AE therapy (BB3RC94 – treatment naïve). (B) Gene expression fold change (FC) between ALDH+ and ALDH− cells of 18 ALDH isoforms detected in the Affymetrix array data. Mean FC for all metastatic samples is represented for each isoform. Red bar indicates isoforms with FC higher than 2. (C) qPCR analysis of *ALDH1A1* and *ALDH1A3* gene expression in the 9 patient metastatic samples that were used in the Affymetrix array. Data is shown as Log10 FC between ALDH+ and ALDH− cells. Mean linear FC of the two ALDH isoforms for all samples is shown. (D) A stably transduced inducible sh*ALDH1A3* MCF-7 cell line was treated with control, tamoxifen (TAM) or fulvestrant (FULV) for 6 days concomitantly with (filled pattern) or without (solid bars) Doxycycline (DOX). *ALDH1A3* mRNA levels were examined by qPCR (left) and percentage of ALDH+ cells was assessed using the Aldefluor assay (right). (E) Venn diagram illustrates meta-analysis of the MCF-7 cell line (control, tamoxifen, fulvestrant-treated ALDH+ vs. ALDH− cells) and the patient Affymetrix data (ALDH+ vs. ALDH− cells). iPathway guide software tool (AdvaitaBio) was used to plot the diagrams. The red dashed-line box indicates the 100 genes that are commonly differentially expressed in ALDH+ cells of patient samples and MCF-7 cell line. The Log2 FC cutoff applied to the ALDH+ vs. ALDH− cells obtained from the meta-analysis data was 0.6. (F) Ingenuity Pathway Analysis (IPA) diagram showing that IL1β signalling is predicted to be activated (orange colour) in the ALDH+ cell population. Straight arrows indicate network of 15 genes, predicted to be regulated by IL1β, that were up-regulated (in red) in ALDH+ cells.

In order to identify which isoforms of ALDH are responsible for the Aldefluor activity of ALDH+ cells in AE resistant ER+ patient samples, we investigated the mRNA expression levels of the 18 detected ALDH isoforms in our patient sample dataset. *ALDH1A1* and *ALDH1A3* showed the greatest fold change (FC) between ALDH+ and ALDH− cells with a mean FC higher than 2 (**Figure 2B**). Validation by qRT-PCR confirmed the elevated expression of *ALDH1A1* and *ALDH1A3* isoforms in the ALDH+ compared to ALDH− population, with a considerably higher averaged linear FC of *ALDH1A3* (300-fold) than *ALDH1A1* (19-fold) across the 9 patient samples (**Figure 2C**). Interestingly, we also found that 6-days of AE treatment significantly up-regulated *ALDH1A3* mRNA levels in two ER+ cell lines, MCF-7 and T47D (**Figure S2A**). Therefore, we used a doxycycline-inducible shRNA system to test the effects of *ALDH1A3* silencing on AE resistance. *ALDH1A3* was stably down-regulated by 58% compared with transfected cells not exposed to doxycycline, and there was a significant decrease in the induction of *ALDH1A3* mRNA levels following AE treatment in the knockdown (KD) cells (**Figure 2D**, left). The enrichment in the ALDH+ cell population after tamoxifen and fulvestrant treatments was significantly reduced in the ALDH1A3KD cells (**Figure 2D**, right), indicating the importance of the ALDH1A3 isoform in ALDH+ cell population after AE therapy.

We also interrogated the gene expression profile of ALDH+ and ALDH− populations from AE-treated MCF-7 cells. The meta-analysis from the patient and cell line microarray datasets (FC≥±1.5 and p≤0.05) revealed 100 genes commonly shared between ALDH+ cells of patient samples and ALDH+ cells of the MCF-7 cell line (**Figure 2E**, **Table S2**). Ingenuity Pathway Analysis (IPA) for these genes predicted activation of eight upstream regulators (z-score ≥2.5), including several cytokines; for example, interleukin-1 beta (IL1β, **Figure S2B**). Of the 100 genes identified in the ALDH+ cell population, 15 were predicted to be regulated by IL1β, and these were all up-regulated in the ALDH+ cells which is consistent with activation of IL1β signalling (**Figure 2F**). One of these genes was interleukin-1 receptor type 1 (IL1R1), which binds and transmits the signal of both IL1α and IL1β.

### AE treatment selects for IL1R1-expressing ALDH+ cells

To study the effects of AE treatment on the ALDH+ population at the single cell level, we analysed the expression of *IL1R1* and *ALDH1A3* in 178 individual ALDH+ cells following tamoxifen or fulvestrant treatment. For this purpose we combined flow cytometry, the single cell C1 system and the Biomark HD technology. Sorted ALDH+ cells were injected and captured in the C1 system, followed by microscopic examination of cell singlets (**Figure 3A**). When comparing *IL1R1* gene expression profiles between control and AE-treated ALDH+ cells, we observed that gene expression levels of *IL1R1* increased significantly following tamoxifen and fulvestrant treatment (**Figures 3B-3C**). In contrast *ALDH1A3* expression is high in nearly all ALDH+ cells, with or without therapy, as expected (**Figures 3B-3C**). *IL1R1* gene expression density plots revealed that control ALDH+ cells match a bimodal distribution with two distinct transcriptomic states: a population that comprises the majority of cells, which show none or very low *IL1R1* gene expression levels, and a small population of cells showing high *IL1R1* levels. However, following AE therapy the vast majority of ALDH+ cells show high *IL1R1* gene expression levels (**Figures 3C** and **S3**). These results reveal the existence of cellular diversity within the ALDH+ population that can be unravelled by single-cell gene expression profiling, and highlight *IL1R1* as an important gene in AE resistant BCSCs. To determine the clinical significance of identifying IL1R1 as facilitating AE resistance, we assessed *IL1R1* gene expression levels in patient breast tumours. Consistent with our cell line data, we found *IL1R1* gene expression levels to be increased in breast tumors following short-term administration of fulvestrant to patients (**Figure 3D**, Patani et al., 2014). In addition, we observed that *IL1R1* expression was significantly increased upon short and long term aromatase inhibitor (AI) treatment compared to baseline levels in four different patient cohorts totalling 404 patients (Ellis et al., 2017; Dunbier et al., 2013; Turnbull et al., 2015) (**Figure 3E**). Notably, we also found that elevated expression of *IL1R1* in ER+ patients who had been treated with AI for 2 weeks was significantly associated with poor outcome (**Figure 3F**).

**Figure 3:**
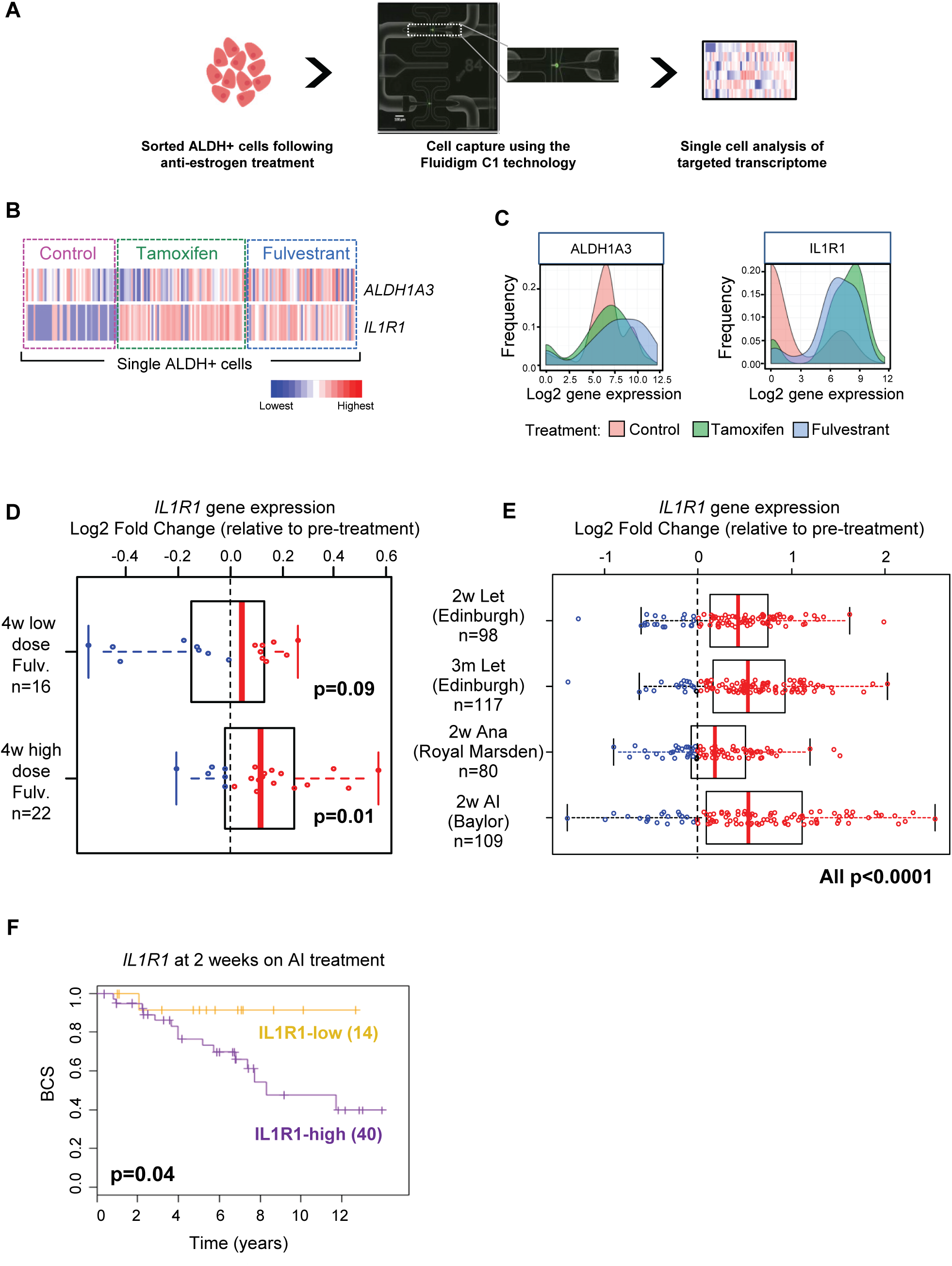
Single ALDH+ cell gene expression in the MCF-7 cell line identifies *IL1R1* overexpression following AE treatment. (A) Schematic overview of the experimental approach to profile single ALDH+ cells. MCF-7 cells treated with either tamoxifen, fulvestrant or control were sorted into single cells and transcription profiles of genes of interest were obtained and analysed as described in the methods. Number of analysed cells: 178. (B) Heatmap of the relative expression across single ALDH+ cells (columns) for *ALDH1A3* and *IL1R1* genes (rows). Cells are ordered by treatment i.e control (left), tamoxifen (middle) and fulvestrant (right). Colours represent expression levels from highest (red) to lowest (blue). (C) Density plots of gene expression in all single ALDH+ cells analysed from the two different AE treatments and control. (D) Box and scatter plot shows *IL1R1* expression from ER+ BC after pre-surgical 4-week treatment with fulvestrant (low-dose, 250 mg, or high-dose, 500 mg) compared to *IL1R1* expression before treatment (Patani et al., 2014). Data is represented as Log2 fold change. Each patient sample is displayed as a blue (down-regulation) or red (up-regulation) circle. P-value calculated with paired Wilcoxon test. (E) Box and scatter plot shows *IL1R1* Log2 fold change gene expression in three different patient cohorts in response to 2 weeks (2w) or 3 months (3m) of letrozole (Let, Edinburgh dataset), anastrozole (Ana, Royal Marsden) and aromatase inhibitor (AI, Baylor dataset) treatment compared to pre-treatment levels. Each patient sample is displayed as a blue (down-regulation) or red (up-regulation) circle. P-value calculated with paired Wilcoxon test. (F) Kaplan-Meier curves represent BC specific-survival (BCS) for *IL1R1*-high and *IL1R1*-low of a cohort of 54 ER+ BC patients (Edinburgh) who received 2 weeks of AI treatment. P-value is based on a log-rank test.

### Single-cell RNA profiling identifies a dormant ALDH+ population that is expanded after AE treatment

*IL1R1* and *ALDH1A3* single-cell gene expression revealed heterogeneity within ALDH+ cells, therefore we decided to investigate the existence of putative subpopulations within the ALDH+ cell population. We examined the mRNA expression level of 68 genes across 444 single ALDH+ MCF-7 cells following control, tamoxifen or fulvestrant treatment. The 68-gene list (**Table S3**) comprised key regulators associated with stemness, self-renewal pathways and markers related to ALDH+ cells that were identified in the whole gene expression data set (Figure 2). A Gaussian Mixture Model approach to estimate and assign clusters to the cells predicted the existence of seven different cellular ALDH+ populations (Control 1 and 2, Tamoxifen 3 and 4, Fulvestrant 5, 6 and 7) (**Figure 4A**). Some of these initial clusters were merged based on their gene expression similarities using Ward’s Hierarchical clustering on Euclidean distance coupled with bootstrapping to estimate branch robustness. This analysis resulted in two major populations of cells, population A and population B which were both made of clusters from the three different treatments, and a small population of Fulvestrant-treated cells (Fulvestrant-7) that were distinct from the rest of the cells (**Figure 4B**). Next, we applied Discriminant Analysis of Principal Components (DAPC) for graphical representation of these three distinct populations (**Figure 4C**). The 8 genes most associated with the first linear discriminant had the highest contribution to the separation of population B from the other populations (**Figure 4D**). Genes associated with cell proliferation, for example the cycle regulator *CCND1* and protein kinase *AKT1*, were downregulated in population B compared to population A, whereas the expression of the mesenchymal marker *SNAI2* was higher in the former (**Figure 4E**). Moreover, population A, which comprised the vast majority of the ALDH+ cells analysed (82%), exhibited higher expression of proliferative markers *PCNA* and *KI67* in comparison to population B (**Figure S4A**). Interestingly, only 10% of the non-treated ALDH+ cells belonged to the quiescent population B, however following tamoxifen and fulvestrant treatment the percentage of quiescent cells represented 44% and 19% of the total cells, respectively (**Figure 4F**). A recent study (Selli et al., 2019) investigated gene expression changes of dormant and acquired resistant ER+ tumours treated with an AI for more than 4 months. Notably, *ALDH1A1* and *ALDH1A3* gene expression levels are significantly increased in dormant tumours compared to acquired resistant tumours which supports the existence of an ALDH+ dormant population after AE treatment (**Figure 4G**). Relative to pre-treatment the dormant tumours also had significantly increased expression of both *ALDH1A1* and *ALDH1A3* as well as *IL1R1* and *SNAI2*, along with reduced *CCND1* (**Figure S4B**) consistent with the results above for the dormant population identified by single-cell analysis. This data suggests that AE resistance can be driven by non-proliferative dormant ALDH+ cells and supports a potential role for IL1R1-targeted therapy to overcome resistance in ER+ BCs (**Figure 4H**).

**Figure 4:**
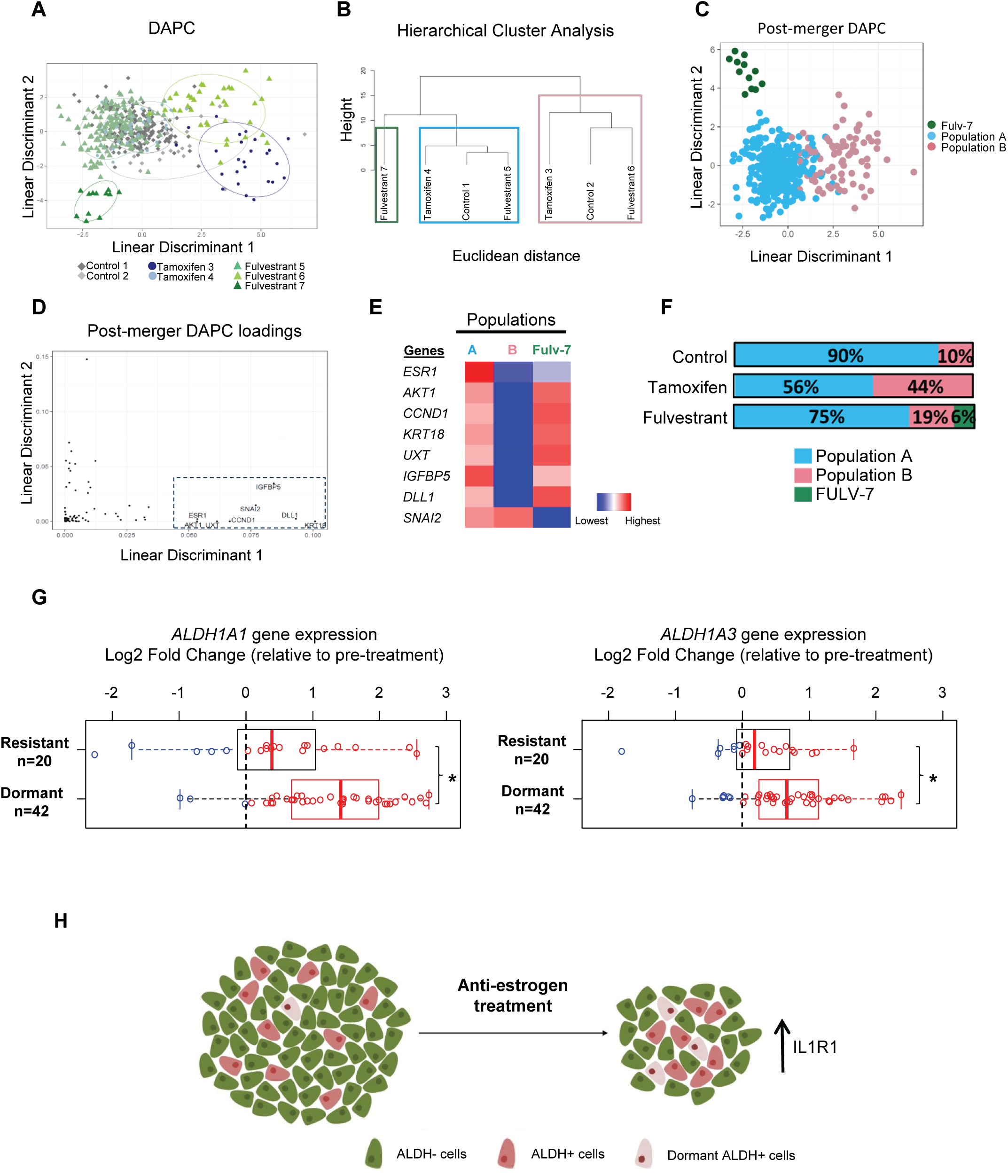
Single-cell gene expression data reveals a dormant ALDH+ population. (A) Scatter plot of the two first linear discriminants from discriminant analysis of principal components (DAPC) analysis for 444 single ALDH+ MCF-7 cells using as classifier the clusters identified through Mclust. The scatter plot shows the cluster of individual ALDH+ cells (rhomboid: control group; circle: Tam group; triangle: Fulv group). Control-treated ALDH+ cells (grey) clustered within two groups (Cluster 1 & 2), Tam-treated ALDH+ cells (blue) also clustered within two groups (Cluster 3 & 4) and Fulv-treated ALDH+ cells (green) clustered within three groups (Cluster 5, 6 & 7). (B) Ward hierarchical clustering of cell clusters using Euclidean distance of all the genes. Boxes represent clusters with an unbiased p-value over 0.90 indicating that these clusters are robust, thus identifying three groups of cells, two major ones, renamed as population A (blue box) and population B (pink box), and a smaller one corresponding to Fulvestrant-7 (green box). (C) Scatter plot of the DAPC analysis for single ALDH+ MCF-7 cells after treatment using as classifier the clusters identified in (B). Linear discriminant 1 accounts for most of the differences between population B and the other two. (D) Distribution of the gene importance to build linear discriminants 1 and 2. Genes above threshold 0.05 of Linear Discriminant 1 are labelled. (E) Heatmap of relative gene expression across the three ALDH+ populations identified (A, B, Fulv-7) for the 8 most important genes in the separation between population B and the others. Colours represent expression levels from highest (red) to lowest (blue). (F) Bar charts show the percentage contribution of each ALDH+ sub-population within the ALDH+ cells treated with control, tamoxifen or fulvestrant. (G) Box and scatter plots show *ALDH1A1* and *ALDH1A3* expression from ER+ dormant and acquired resistant tumours after 4-months neoadjuvant treatment with letrozole compared to expression before treatment (Selli et al., 2019). Data is represented as Log2 fold change. Each patient sample is displayed as a blue (down-regulation) or red (up-regulation) circle. P-value calculated with paired Wilcoxon test. (H) Diagram showing that AE therapies do not target ALDH+ cells and enrich for a dormant IL1R1+ALDH+ cell population.

## DISCUSSION

Previously, we reported that ALDH+ cells are resistant to AE therapy and that high ALDH1 expression predicts resistance in women treated with tamoxifen (Simões et al., 2015). Our findings here establish a role for the IL1R1 signalling pathway in the regulation of AE resistant ALDH+ BCSCs. We also identify heterogeneity in the ALDH+ cell population and an expansion of a quiescent ALDH+ sub-population after AE therapies.

Firstly, we showed that BC cells treated with AEs contain a population of ALDH+ cells that have higher mammosphere-forming and tumour-initiating cell frequency than ALDH− cells. We next wanted to further characterize these cells and the mechanisms that drive them. ALDH1A1 and ALDH1A3 isoforms are both reported to be predictive biomarkers of poor clinical outcome in BC (Liu et al., 2014; Marcato et al., 2011), and we found them to be the most highly increased among 18 ALDH isoforms detected in metastatic patient-derived ALDH+ BC cells. ALDH1A3 knock down confirmed that this isoform is crucial for enriching the ALDH+ population following AE treatment. This data supports the growing body of literature describing the involvement of ALDH1A3 in cancer stemness, tumour progression and poor prognosis.

We found that ALDH+ cells have a different gene expression profile compared to ALDH− cells in both ER+ metastatic patient samples and MCF-7 cells. In particular, genes that predicted activation of pro-inflammatory cytokine IL1β signalling, including *IL1R1*, were expressed at higher levels in ALDH+ cells. By using single-cell gene expression profiling in the ALDH+ cell population we identified *IL1R1* to be significantly up-regulated in AE resistant ALDH+ cells compared to control cells. Importantly, we found that expression of *IL1R1* is induced in the tumours of patients treated with AE therapies and predicts for treatment failure. This data indicates that IL1β signalling is likely to be important for CSCs to drive AE resistance in BC. *IL1β* expression correlates with increased aggressiveness and enhanced metastatic potential of BC cells, suggesting IL1β as a potential biomarker for predicting which patients are likely to be diagnosed with BC metastasis, specifically to bone (Tulotta et al., 2019).

Single-cell targeted transcriptome analysis revealed the existence of distinct clusters within ALDH+ cells and the expansion of a quiescent ALDH+ population (Population B) after AE therapies. Heterogeneity within BCSCs of MCF-7 cells has previously been described using different CSC functional assays, such as mammospheres, growth in hypoxia and PKH26 retention, to isolate single cells for gene expression analysis (Akrap et al., 2016). Our data suggests that Population B represents a small population of non-dividing quiescent ALDH+ cells that survive AE treatments, which may enable them to survive for long periods of time and eventually lead to late recurrence in ER+ BC patients. This idea is supported by data from AI-induced dormant tumours that express increased levels of *ALDH1A1* and *ALDH1A3* genes. Recently, single-cell RNA profiling of normal breast samples identified four cell clusters within the ALDH+ cell population (Colacino et al., 2018). Interestingly, Population B resembles cluster 3 identified in this publication, which was characterized by high expression of mesenchymal markers, including *SNAI2*, and low expression of proliferative genes, such as *KI67*, *PCNA*, *CCND1*. Population B expresses low levels of *AKT1* and AKT1^low^ cancer cells have been reported to be quiescent cells that survive chemotherapy in breast tumours (Kabraji et al., 2017).

Combination therapies targeting both bulk tumour cells and BCSCs should reduce the probability of tumour relapse and, therefore, pharmacological inhibitors that target CSC pathways have been highly pursued and are being tested in patients (Brooks et al., 2015). In our model, both proliferative and dormant AE resistant BCSCs express IL1R1. This suggests that anti-IL1R1 therapies, such as Anakinra (recombinant form of human IL-1 receptor antagonist) or Canakinumab (human anti-IL1β monoclonal antibody), could represent a new strategy to target AE-resistant CSCs.

In conclusion, the present work contributes to our understanding of the cellular heterogeneity present in the AE resistant BCSC population. Our work suggests that CSC dormancy is an adaptive strategy to evade AE treatments and supports targeting of ALDH+IL1R1+ cells to reverse AE resistance. This work highlights the advantages of single-cell transcriptomic analysis, rather than bulk tissue, to interrogate the cellular heterogeneity within the ALDH+ CSC population. Further understanding of the dormant ALDH+ population that survives AE therapies, particularly using clinical samples, will provide new insights for prevention and treatment of recurrences of ER+ BC.

## EXPERIMENTAL PROCEDURES

A comprehensive description of the methodology is included in the supplementary information.

### Breast Cancer Samples

Consented, de-identified pleural effusion or ascitic fluids were collected at the Christie NHS Foundation Trust (UK) or the University of Michigan (USA). The clinico-pathological details of the samples are shown in **Table S4**.

### ALDH+/− Cell Isolation

BC cells were stained using the Aldefluor assay (Stemcell Technologies) following manufacturer‘s protocol and isolated using the Influx cell sorter (BD Biosciences).

### Single-cell capture and transcriptomics profiling

Single ALDH+ MCF-7 cells were captured within the C1 system using the medium C1 Single-Cell Preamp Integrated Fluidic Circuit (IFC, 10-17 µm) chips (Fluidigm, 100-5480). Individual cells were visualised using the Leica Widefield Low Light microscope. Cell loading, lysis, reverse transcription and cDNA pre-amplification were performed within the C1 system following manufacturer’s instructions. Single-cell transcriptomics profiling of 444 single cells was performed using the 96.96 Dynamic Array IFC Biomark chips (Biomark HD Real-Time PCR System, Fluidigm) to interrogate the expression of 68 TaqMan assays in each cell. Single-cell data was processed and analysed as described in the data analysis section. Two independent experiments with technical replicates were performed.

## AUTHOR CONTRIBUTIONS

B.M.S., R.B.C. conceptualised the study. A.S.-C., R.B.C., B.M.S. designed, carried out the experiments, performed data interpretation and wrote the manuscript. E.C.-G. carried out bioinformatics single-cell analysis and wrote parts of manuscript. A.H.S. performed bioinformatics analysis on Affymetrix data and patient data. M.J. created knockdown cell line. C.C., M.D.B. helped carry out single-cell experiments. M.E.B., D.F.H. provided patient samples from Michigan cohort. A.S.-G., R.E., M.S.W., S.J.H. advised on experimental design and revised the manuscript. All authors edited and approved the final version.

## ACKNOWLEDGEMENTS

We are grateful to Breast Cancer Now (MAN-Q2), Breast Cancer Research Foundation, and the National Cancer Institute (R35 CA197585) for funding this research. B.M.S., R.B.C. and S.J.H. are supported by the NIHR Manchester Biomedical Research Centre (IS-BRC-1215-20007). We would like to thank Medical Research Council that funded A.S-C with a Doctoral Training Scholarship (MR/K501311/1). We are grateful to the European Association of Cancer Research and the Windgate Foundation for funding travel fellowships to A.S-C. M.S.W. has financial holdings in Oncomed Pharmaceuticals and receives research support from MedImmune.

---

**Figure S1.**
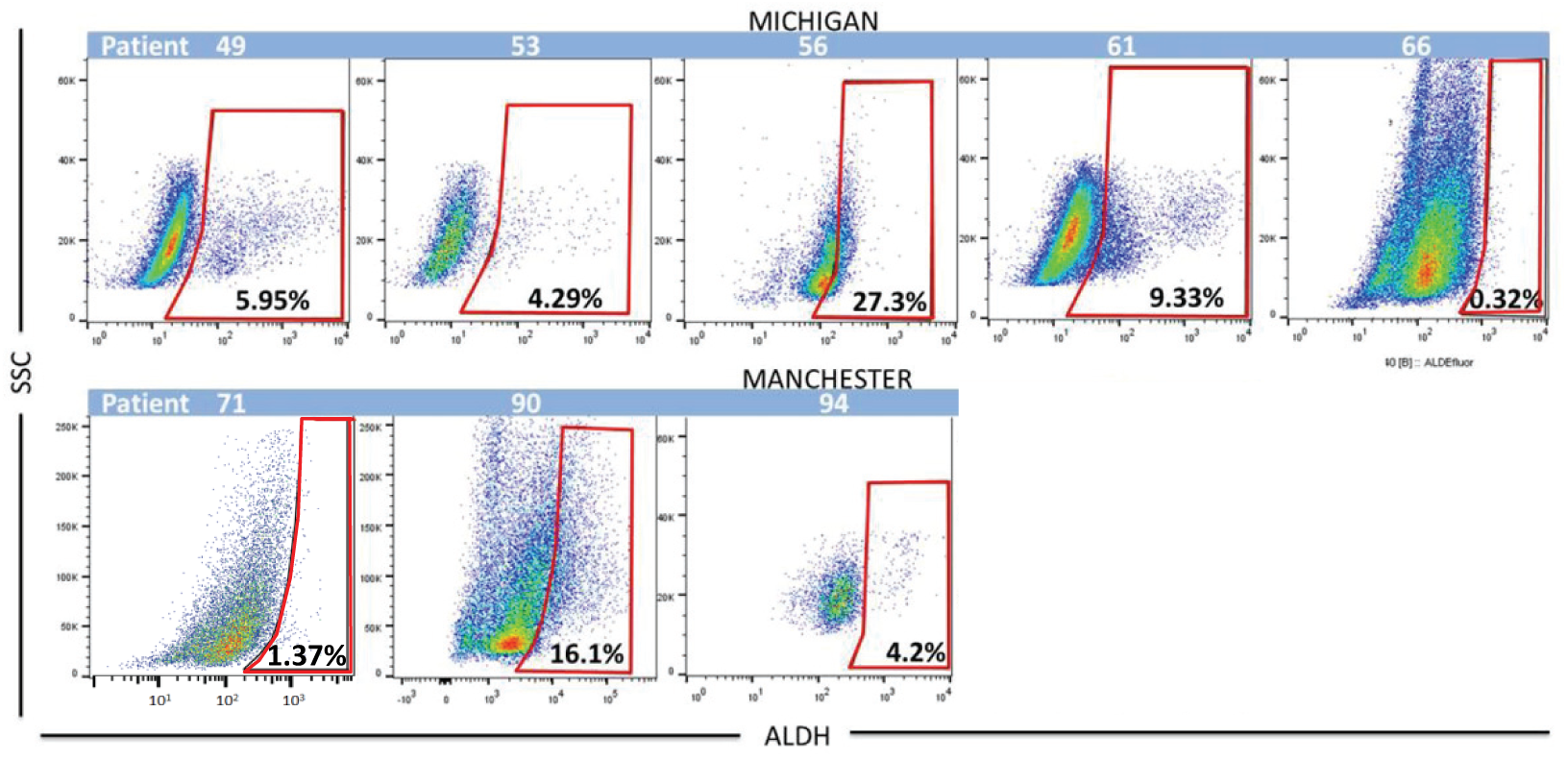
FACS plots showing percentage of ALDH pos cells, measured by the Aldefluor assay, in metastatic patient samples. ALDH pos cells (red box) from Michigan‘s biobank (top) and Manchester‘s biobank (bottom) patient-derived samples are shown.

**Figure S2.**
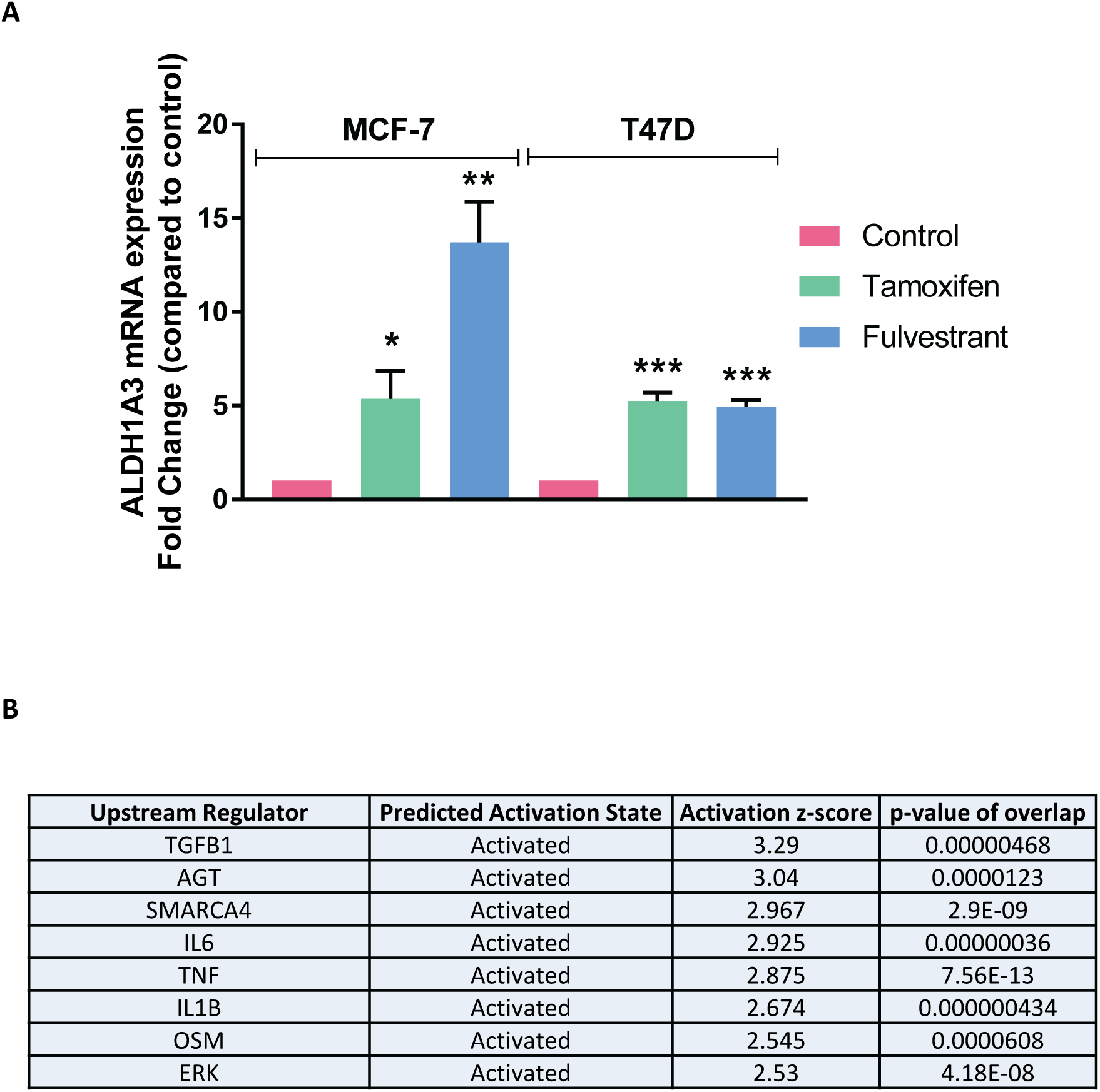
**A)** ALDH1A3 mRNA expression in MCF-7 and T47D cells following tamoxifen (green) and fulvestrant (blue) treatment compared to control (pink). **B)** List of upstream regulators and respective predicted activation (with z-score ≥2.5) identified by Ingenuity Pathway Analysis (IPA) of 100 genes commonly expressed in ALDH+ cells of patient samples and MCF-7 cells.

**Figure S3.**
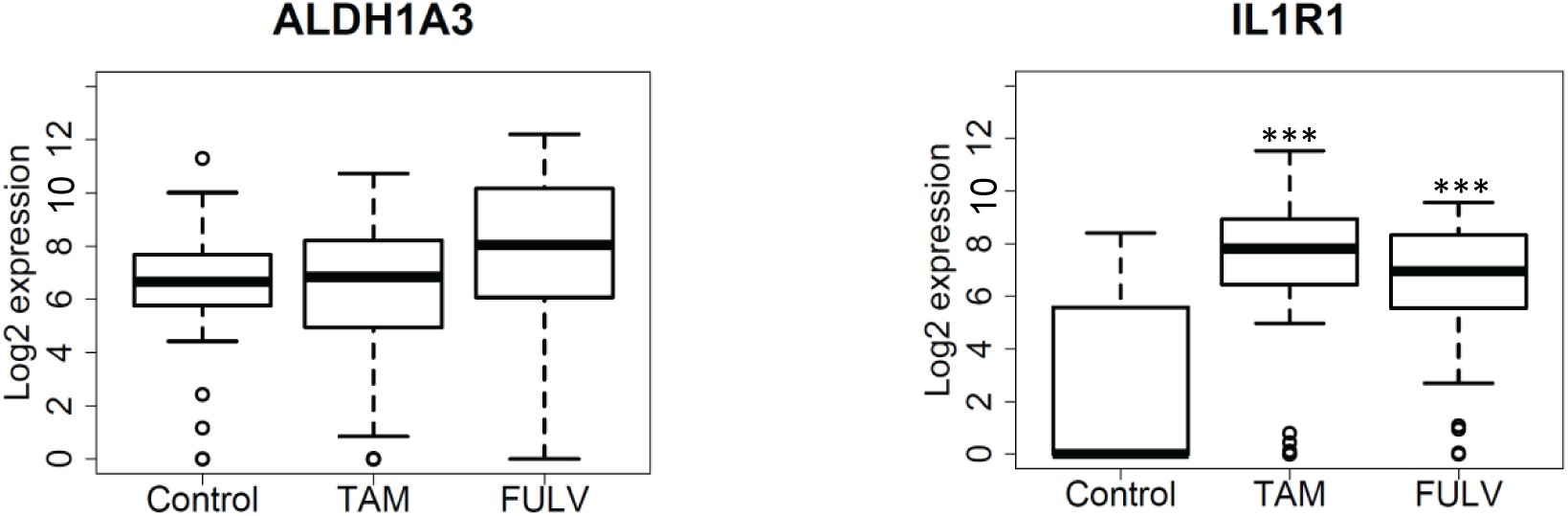
Boxplots of *ALDH1A3* and *IL1R1* gene expression in cells treated with vehicle (Control), tamoxifen (TAM) and fulvestrant (FULV). Log2 expression distribution in all the cells is shown as boxes containing the interquartile ratio (first and third quartiles) with the median (bold line) and whiskers representing the 5–95% range. Kruskal-Wallis with Dunn’s post-hoc correction was used to compare tamoxifen/fulvestrant treated cells versus control cells.***Pvalue<0.001

**Figure S4.**
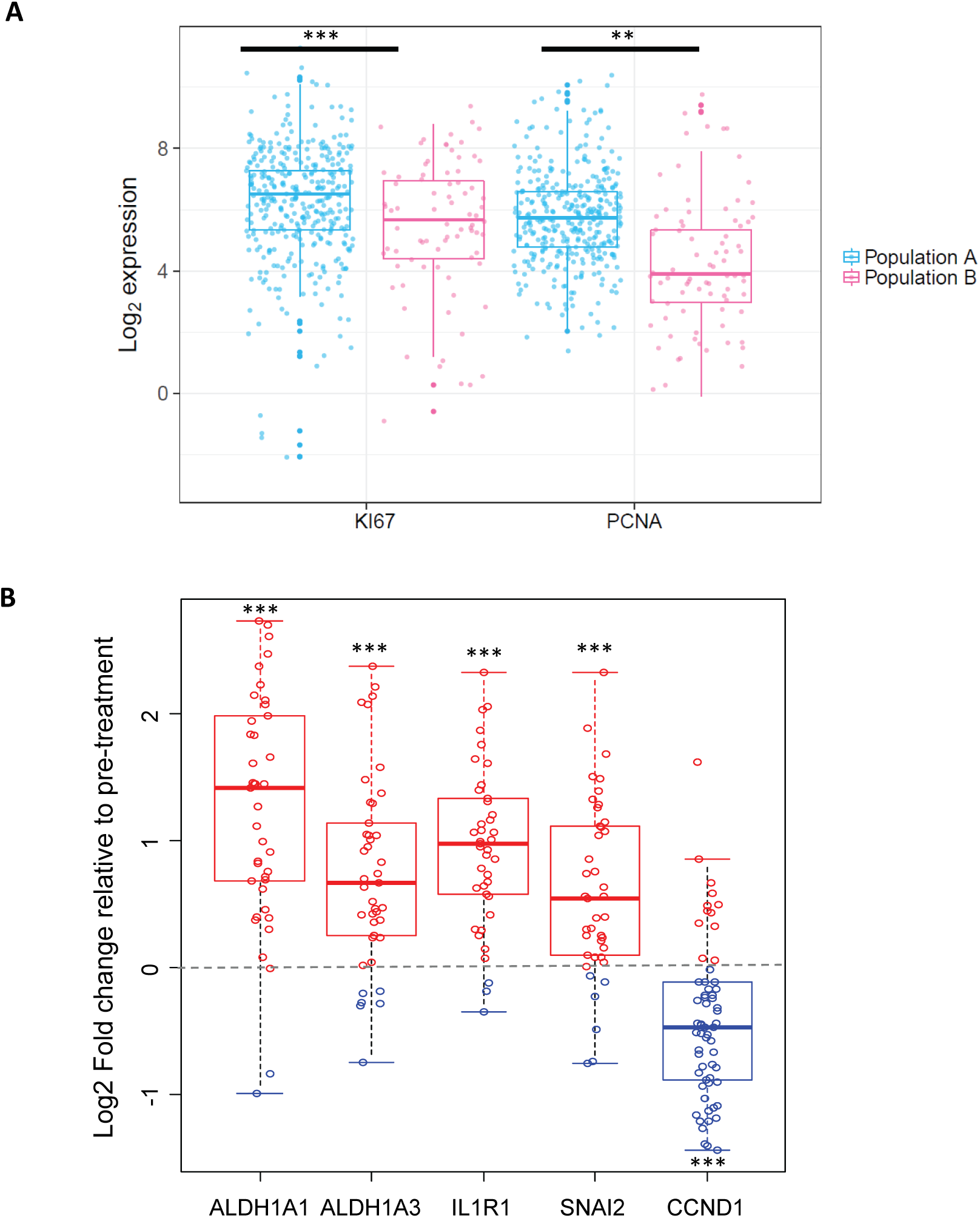
**A)** Boxplots of *Ki67* and *PCNA* gene expression in cells of Population A and B. Log2 expression distribution in all the cells is shown as boxes containing the interquartile ratio (first and third quartiles) with the median (bold line) and whiskers representing the 5–95% range. Each point within the plot represents a cell for the gene specified. Mann-Whitney U Test was used to compare Population B vs Population A. P values were adjusted for FDR with Benjamini and Hochberg method. (***Adj -Pvalue <0.001; **Adj-Pvalue<0.01). **B)** Boxplots show *ALDH1A1*, *ALDH1A3*, *IL1R1*, *SNAI2, CCND1* gene expression from ER+ dormant tumours after 4-months neoadjuvant treatment with letrozole compared to expression before treatment (Selli et al., 2019). Data is represented as Log2 fold change. Each patient sample is displayed as a blue (down-regulation) or red (up-regulation) circle. P-value calculated with paired Wilcoxon test.

## Supplementary Tables

**Table S1.** Please refer to Supplemental spreadsheet file.

**Table S2.** Please refer to Supplemental spreadsheet file.

**Table S3.** List of 68 genes used for single-cell targeted transcriptome analysis.

| Genes |  |  |  |
| --- | --- | --- | --- |
| *ABCG2* | *EPCAM* | *ITGA6* | *NUMB* |
| *AKT1* | *ESR1* | *JAG1* | *PCNA* |
| *ALDH1A3* | *EZH2* | *JAG2* | *PGR* |
| *AR* | *FBXW7* | *KRT18* | *PIK3CA* |
| *BRCA1* | *GAPDH* | *KRT19* | *POU5F1* |
| *CA12* | *GATA3* | *KRT8* | *RAB7A* |
| *CCND1* | *GJA1* | *LIN28A* | *SNAI2* |
| *CD24* | *GPRC5A* | *MET* | *SOCS3* |
| *CD44* | *GSK3B* | *MKI67* | *SOX2* |
| *CDH1* | *HER2* | *MKP1* | *TAZ* |
| *CDH3* | *HES1* | *MTOR* | *TGFB1* |
| *CTNNB1* | *HPRT1* | *MUC1* | *TGFBR1* |
| *CXCR1* | *ID1* | *NANOG* | *TP53* |
| *CXCR4* | *IGFBP5* | *NFKB1* | *TSPAN6* |
| *CYR61* | *IL1R1* | *NOTCH1* | *TWIST1* |
| *DLL1* | *IL6R* | *NOTCH2* | *UXT* |
| *ENAH* | *IL6ST* | *NOTCH3* | *YAP1* |

**Table S4.** Clinico-pathological information of the Michigan and Manchester patient datasets. ND: No data available; QS: Quick score.

| Patient | ER | PR | Chemotherapy | Endocrine Therapy | Bone Therapy | Other Therapy | Metastasis |
| --- | --- | --- | --- | --- | --- | --- | --- |
| **Mi49** | + | + | Capecitabine<br>Cyclophosphamide<br>Doxorubicin<br>Eribulin<br>Etoposide<br>Gemcitabine<br>Ixabepilone<br>Taxol | Anastrozole<br>Fulvestrant<br>Goserelin | Zometa<br>Zoledronic acid | Anti-DLL4<br>antibody:<br>OMP-21M18 | Bone<br>Brain<br>Pleura |
| **Mi53** | + | + | Cyclophosphamide<br>Doxorubicin<br>Taxol | Anastrozole<br>Tamoxifen | Zometa | ND | Bone<br>Liver |
| **Mi56** | + | + | Capecitabine<br>Carboplatinum<br>Paclitaxel<br>Vinorelbine | Exemestane<br>Letrozole<br>Tamoxifen | Denosumab | ND | Bone<br>Chest wall<br>Lymph node |
| **Mi61** | + | + | Cyclophosphamide<br>Docetaxel<br>Doxorubicin | Anastrozole<br>Exemestane | Denosumab | ND | Bone<br>Pleura |
| **Mi66** | + | + | Capecitabine<br>Carboplatin<br>Gemcitabine | Arimidex<br>Tamoxifen | Denosumab | ND |  |
| **BB3RC68** | QS8 | ND | Capecitabine<br>Fluorouracil + Epirubicin +<br>Cyclophosphamide (FEC) | Anastrozole<br>Fulvestrant<br>Tam | ND | ND | Bladder<br>Liver<br>Lung<br>Lymph nodes<br>Peritoneum |
| **BB3RC69** | 96% | 77% | ND | Anastrozole<br>Letrozole<br>Tam<br>Fulv | Pamidronate<br>Zoledronic Acid | ND | Bone<br>Lymph node<br>Peritoneum |
| **BB3RC71** | 54% | 72% | Capecitabine<br>Eribulin<br>FEC<br>Taxol<br>Taxotere<br>Vinorelbine | Anastrozole<br>Exemestane<br>Fulvestrant<br>Goserelin<br>Letrozole<br>Tamoxifen | Pamidronate<br>Zoledronic Acid<br>Ibandronic Acid | Herceptin<br>Lapatinib | ND |
| **BB3RC89** | QS 8 | QS 8 | Capecitabine<br>FEC<br>Taxol | Letrozole<br>Tamoxifen | ND | ND | Bone<br>Liver<br>Lung |
| **BB3RC90-90A** | QS 8 | QS 8 | Capecitabine | ND | ND | ND | Bone<br>Liver<br>Pleura |
| **BB3RC91-91A** | 96% | 98% | Docetaxel<br>FEC | Anastrozole<br>Exemestane<br>Letrozole<br>Tamoxifen | ND | Everolimus | Bone<br>Liver<br>Omentum<br>Peritoneum |
| **BB3RC94** | + | + |  | Treatment naïve |  |  | ND |

N.B. Samples 90 and 90A are from the same patient but were taken at different time points. The same applies to samples 91 and 91A.

## SUPPLEMENTAL EXPERIMENTAL PROCEDURES

### Breast Cancer samples

Metastatic fluids were collected at the Christie NHS Foundation Trust (UK) in accordance with local research ethics committee guidelines (study number: 05/Q1402/25) or the University of Michigan (study number: IRBMED 2001-0344/HUM00042204). Fluids were spun at 1000 g for 10 min at 4°C and pellets were resuspended in Phosphate Buffered Saline (PBS). Erythrocytes and leucocytes were depleted from the metastatic fluids by using density gradient Lymphoprep (Stemcell Technologies) following manufacturer‘s protocol. Clinical information about patient samples is shown in **Table S4**.

### ALDH+/− Cell Isolation

Breast cancer cells were re-suspended in Aldefluor buffer and incubated in the presence of the Aldefluor reagent bodipyaminoacetaldehyde (BAAA) (Aldefluor assay, Stemcell Technologies) for 40 minutes at 37°C, following the manufacturer‘s protocol. A subset of cells were also incubated with the selective ALDH inhibitor diethylaminobenzaldehyde (DEAB) to distinguish between ALDH+ and ALDH− cells. When performing single-cell experiments using the C1 system (Fluidigm), cells were stained for CD44 (CD44-APC; BD,) and CD24 (CD24-PECY7) expression as well as ALDH activity in order to isolate ALDH+ cells that are not CD44^high^ CD24^low^. Following incubation, cells were washed with PBS and stained with the cell viability dye 7-aminoactinomycin (7AAD, BD). Cells were then FACS-sorted into 200 µl of 2% Fetal Bovine Serum in Hank’s Balanced Salt Solution (HBSS) using the InFlux (BD bioscience). Single colour stains were included for compensation and gating purposes. Data was analysed using FlowJo 10.1.

### Mammosphere Culture

Cells were counted and seeded at a density of 5,000 cells/well in 6-well plates containing mammosphere media (DMEM/F12 media with L-Glutamine (Gibco), B27 supplement (Gibco; 12587) and 20 ng/ml EGF (Sigma)) for 7 days at 37°C. Mammosphere forming efficiency (MFE) was calculated by dividing the number of mammospheres greater than 50µm by the number of cells seeded per well and is expressed as the mean percentage of MFE (Shaw et al., 2012).

### Transplantation Assays

MCF-7 cells were treated *in vitro* with 10^−6^ M 4-Hydroxytamoxifen (Sigma-Aldrich, H7904), 10^−7^ M fulvestrant (ICI 182,780, Tocris, 1047) or ethanol (vehicle) for 6 days following staining with the Aldefluor assay. Serial limiting dilution of sorted ALDH+ and ALDH− cells (10,000; 1,000; 100 cells) were resuspended in mammosphere media mixed 1:1 with Matrigel (BD bioscience, 356234) and inoculated subcutaneously into the left and right flanks of female NOD/SCID Gamma (NSG) mice. All in vivo work was carried out using n=4 mice for each condition. 90-day slow release estrogen pellets were implanted subcutaneously into mice before cell injection (0.72 mg, Innovative Research of America) and, after day 90, 8 μg/ml of 17-beta estradiol was administered in drinking water. Tumour measurements were taken three times a week and tumour size was calculated using the formula:

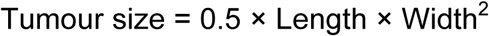

Positive tumour growth was assessed at week 20 after cell injection by determining the mice bearing a tumour greater than 300 mm^3^. Extreme Limiting Dilution Analysis (ELDA) was performed using software available at http://bioinf.wehi.edu.au/software/elda/ (The Walter and Eliza Hall Institute of Medical Research) to assess differences in stem cell frequency.

### RNA extraction and Real-Time PCR

ALDH+ bulk cells (≥10,000 cells) were sorted into 100 µl of lysis buffer containing 1% β-Mercaptoethanol, following by cell disruption and homogenisation via vortexing for 1 minute. RNA was extracted using the RNeasy Plus Micro Kit (Qiagen, 74034) with on-column DNase treatment following manufacturer’s protocol. The Bioanalyzer (Agilent 2100 Bioanalyzer system, Agilent Technologies) and the Qubit (Thermofisher Scientific) were used for quantitation and quality control of the RNA.

### Bulk transcriptome analysis

Human Array Gene 1.0 ST (Affymetrix) GeneChips were used to assess mRNA expression profile in bulk ALDH+ and ALDH− cells. Double stranded amplified cDNA was generated using the Ovation Pico WTA System V2 (NuGen) and the Single Primer Isothermal Amplification (SPIA) following manufacturer’s guidelines. cDNA was fragmented and labelled prior hybridisation onto the array (GeneChip hybridization Oven 640, Affymetrix). The GeneChip array was then washed and stained using the Fluidics Station protocol FS450_0007 and the Affymetrix GeneChip Command Console Software (Affymetrix) following manufacturer‘s guidelines. The GeneChip array was scanned using the Scanner 3000 system with autoloader (Affymetrix).

Microarray data from cell line and patient samples were processed using the *Affy* package in R (Gautier et al., 2004). Data was quantile-normalised and Log2 transformed. Differential gene expression analysis was carried using paired Rank Products (Breitling et al., 2004). Meta-analysis was performed using iPathwayGuide (AdvaitaBio). Statistical significance for RNA expression was assessed using t-test parametric testing.

### Single-Cell Analysis

Data generated by the Biomark (Fluidigm) were converted into Log2 expression values and quality controls were undertaken. These included data filtering to remove all values under the limit of detection, which was set to threshold cycles (Ct) greater than 28; the removal of genes expressed in 3 or less cells within each treatment and, the removal of outliers (via the function *identifyOutliers* implemented in the R package FluidigmSC - Fluidigm Corporation, 2014). Missing completely at random values were estimated and inputted using the R package MICE (Azur et al., 2011). We undertook a statistical approach to eliminate doublets derived from equipment unfitness (Fluidigm Corporation, 2016). Using the package Mclust in R we fitted Gaussian mixture models to identify cell clusters within each treatment. These models indicated the existence of two very well defined cell clusters in each condition. The nature of these clusters was further investigated by plotting the average Log2 expression per gene in both clusters, pointing towards a stratification into doublets and singlets. Clusters corresponding to singlets were taken forward for the analysis. With the aim of identifying different cell populations we used a finite Gaussian mixture model to (a) estimate the number of clusters within the data (function Mclust within the R package Mclust (Scrucca et al., 2016)) and (b) generate those clusters. Ward hierarchical clustering with bootstrapping (Ward, 1963) was undertaken with the package pvclust in R to find similarities between clusters and merge smaller clusters into larger ones (threshold of Approximately Unbiased (AU) p-value greater than 0.9). Merged clusters were assessed for biases regarding batch and plate. Further analysis of cluster similarities and genes associated to cluster differences were undertaken using Discriminant Analysis of Principal Components (DAPC) (Jombart et al., 2010) of the 7 clusters identified with Mclust and fitting a model built using 40 principal components (PCs) and 8 linear discriminants. The number of PCs to use to build the model was determined via cross-validation by building 1000 different models per PC with an 80-20 split of training/test data and selecting the combination that provided the maximum correct predictions with the lowest number of PCs. Further merging of the clusters was further assessed using DAPC to find the differences between the three main populations of cells identified (A, B and Fulv7) and built with 40 PCs and 6 linear discriminants. Selected genes were compared across conditions using non-parametric Mann-Whitney test. False discovery rate was controlled via Benjamini and Hogberg method.

### shRNA knockdown

The inducible Dharmacon TRIPZ lentiviral shRNA was used to stably down-regulate ALDH1A3 mRNA expression levels (ALDH1A3KD - Dharmacon, V3THS_378581; V3THS_378584; V3THS_378585).

### Statistical Analysis

P values less that 0.05 were considered significant (*p<0.05, **p<0.01, ***p<0.001). Results are presented as the mean of at least 3 independent experiments ± Standard Error of the Mean (SEM) or Standard Deviation (SD).

### Accession Numbers

The Affymetrix data in this paper have been deposited in NCBI’s Gene Expression Omnibus repository under series accession number GSE136287.

